# Environmental DNA: a new low-cost monitoring tool for pathogens in salmonid aquaculture

**DOI:** 10.1101/215483

**Authors:** Lucy Peters, Sofie Spatharis, Maria Augusta Dario, Inaki J T Roca, Anna Kintner, Øyvind Kanstad-Hanssen, Martin S. Llewellyn, Kim Praebel

## Abstract

Sequencing of environmental DNA (eDNA-seq) is an emergent new monitoring tool that promises to facilitate the accurate and cost effective detection of species in environmental samples. eDNA monitoring is likely to have a major impact on the ability of salmonid aquaculture industry producers and their regulators to detect the presence and abundance of pathogens and other biological threats in the surrounding environment. However, for eDNA-seq to develop into a useful bio-monitoring tool it is necessary to (a) validate that sequence datasets derived from amplification of meta-barcoding markers reflect the true species’ identity and abundances in biological samples, and (b) establish a low-cost sequencing method to enable the bulk processing of environmental samples. In this study, we employed an elaborate experimental design whereby different combinations of five biological agents were crossed at three abundance levels and exposed to pre-filtered and normal seawater, prior to coarse filtering and then eDNA ultrafiltration of the resultant material. We then benchmarked the low-cost, scalable, Ion Torrent sequencing method against the current gold-standard Illumina platform for eDNAseq detection in aquaculture. Based on amplicon-seq of the 18S SSU rDNA v9 region, we found that Illumina and Ion Torrent were equally good in identifying the two parasite species (*Lepeophtheirus salmonis* and *Paramoeba perurans),* whereas the microalgae species *Prymnesium parvum*, *Pseudo-nitzschia seriata* and *P. delicatissima* could be assigned correctly only to the genus level. Illumina and Ion Torrent were also equally able to reflect community composition in our samples, whereas Ion Torrent was more sensitive in detecting species richness when the medium was unfiltered seawater. Both methods were able to reflect the correct abundances of 4 out of 5 species in samples from unfiltered seawater, despite the significant amount of background noise from both bacteria and eukaryotes. Our findings indicate that eDNA-seq offers significant potential in the monitoring of species harmful to aquaculture and for this purpose, the low-cost Ion Torrent sequencing is equally as accurate as Illumina.

## Introduction

The salmonid aquaculture industry is undergoing explosive growth globally. However, the industry is beset by parasitic disease and is often the subject to mass mortalities of farmed fish due to toxin-producing Harmful Algal Blooms (HABs) ^1^. Economic losses associated with certain agents, as for example sea lice, accounts for up to £ 470 M / year for major producers like Norway ^2^. The presence and abundance of these potentially damaging organisms in the environment and around aquaculture sites must be closely monitored. Traditional microscopy methods for algal and copepod larval species identification and abundance estimates are time-consuming, demand expertise and are not always accurate when abundances are low or when cryptic species are involved. Similarly, parasite counts on the fish themselves are both time consuming and impose significant handling stress. Environmental DNA (eDNA) analysis is an emerging molecular approach for species identification from samples containing cellular DNA and extracellular DNA sloughed off all living organisms ^3^. eDNA analysis has been successfully employed to detect and monitor eukaryotic micro- and microbial communities and populations ^4^ and is a useful tool for early monitoring systems as it allows for more accurate and standardized detection of species that are cryptic, inaccessible ^4b^ and of low abundance ^4a^. Some recent advances have been made in its application for pathogen detection in freshwater aquaculture ^5^. However, before it can be considered as a systematic bio-monitoring tool it is necessary to find a cost-effective genotyping approach to allow rapid processing of environmental samples on a day-to-day basis. Furthermore, validation is required to establish a quantitative relationship between eDNA genotype data and biological abundance (e.g. ^6^).

Next generation sequencing (NGS) methods are increasingly being used to characterize and monitor aquatic organisms from eDNA samples ^7^. NGS can also be used for metabarcoding – the use of universal primers to amplify DNA from many different organisms within one sample. Moreover, barcoding of sequences within the same sample allows for parallel processing of multiple samples both during the sequencing run as well as during the downstream bioinformatic analysis ^8^. Despite the relatively high cost of NGS, cost-efficiency choices between different platforms can be made based on suitable benchmarking.

In this study, our aim was to establish whether amplicon sequencing can reflect the known abundance and diversity of eukaryotic aquaculture pathogens and harmful algae. Our first objective was to use an elaborate experimental approach involving cross treatments of the aforementioned aquaculture pathogens controlling for background noise influence (i.e. unfiltered sea-water) versus filtered sea-water. We then benchmarked Illumina MiSeq and Ion Torrent techniques to deep sequence the ribosomal 18rRNA marker gene of these samples. Our final objective was to compare the sequencing techniques in terms of their sensitivity to detect species and reflect their abundances, and to discriminate between treatments based on the sample diversity and whole community composition.

## Methods

### Choice of pathogen

Four major pathogens and risk agents were selected: *Lepeophtheirus salmonis* and *Paramoeba perurans* as well as the algal risk agents *Prymnesium parvum* (identified morphologically in University of the Aegean)*, Pseudo-nitzschia seriata* (identified by N. Lundholm personal communication), and *P. delicatissima* (CCAP culture). *L. salmonis* is one of the most important and widespread, affecting farmed Atlantic salmon in Norway, Ireland and the UK^9^. *N. perurans* is the causative agent of amoebic gill disease, which is a major source of commercial loss for aquaculture in Tasmania, but also affects industries in both North and South America as well as in Europe ^10^. The haptophyte *P. parvum,* produces compounds known as prymnesins causing severe toxic effects by affecting plasma membrane integrity of gills ^1b, 11^, Oikonomou et al., 2012). *P. parvum* blooms resulted in extensive mortality of farmed salmon in Scotland and Norway in the past ^1b^. *Pseudo-nitzschia* produce the neurotoxin Domoic acid (DA), that causes Amnesic Shellfish Poisoning (ASP) symptoms in humans ^1a^ when it bioaccumulates in the tissue of bivalves.

### Incubations and filtering

Four different groups of incubations were set up, containing a single species or combinations of the pathogens. Each of the four groups was further divided into three treatment regimens of the following abundances: two, six and eighteen, in the case of the salmon louse referring to number of individuals (females without egg strings), and to cell counts for the amoeba and algal species. The densities of the cultures of amoeba and algal species were determined by microscopy and dilution series in a Neubauer chamber. As a negative control, a blank baseline treatment was set up for each of the two different days of the incubations. All incubations consisted of 2.0 L of two different media; sterile filtered (0.22μm) seawater and unfiltered seawater. In this way, the strength of the relationship between known abundances and sequence read numbers could be tested both for the baseline level of PCR amplification only, as simulated by the filtered medium, and in the context of the natural environment, replicated by the unfiltered medium. Both the unfiltered and filtered seawater were obtained from internal pipes of the marine laboratory of the Norwegian College of Fishery Science, UiT the Arctic University of Norway. Our experimental design generated 72 samples (2 medium types: filtered/unfiltered x 12 treatments x 3 replicates) (see Figure 1 for experimental design). For each medium type, we also used 2 blanks replicated thrice (12 samples in total), resulting in a total of 84 samples. All incubations were performed at 3.8°C for 24 hours, before 0.5 L were filtered to collect material for DNA extraction and sequencing through a 0.22µm sterivex filter unit (EMD Millipore, Darmstadt, Germany). Samples for all incubations were obtained in triplicate.

**Figure 1.**
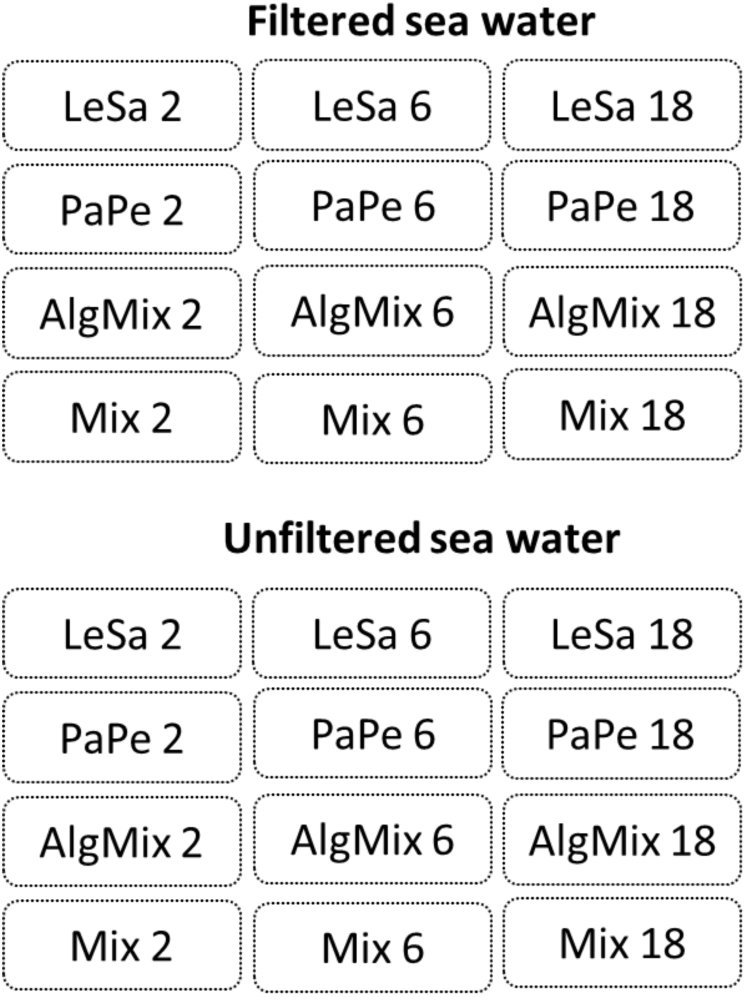
Experimental design aiming to test the efficiency of two sequencing methods, namely Illumina MiSeq and IonTorrent, in reflecting the species composition and abundance in different incubations. Each incubation contained four aquaculture related risk agents either in isolation or combined: *Lepeophtheirus salmonis* (LeSa), *Paramoeba perurans* (PaPe), algal mix (AlgMix) containing *Prymnesium parvum*, *Pseudonitzschia delicaticima and P. seriata,* and Mix containing all the species together. Each of these incubations was performed at three abundance levels representing triplings of the initial abundance (i.e. 2, 6 and 18). The experiment aimed to control for the effect of background noise in these treatments thus we deployed these using both filtered and unfiltered seawater medium.

### DNA barcoding and sequencing

Total DNA was extracted using the DNeasy Blood and Tissue kit (Qiagen, Hilden, Germany) following the manufacturer’s instructions with the minor adjustments. Briefly, 500µl instead of 200 µl digestion buffer ATL/proteinase K was added directly to each Sterivex filter and incubated over night with continues rotation. The buffer, containing the eDNA, was then spun out of the filters into a 2 ml Eppendorf tube at 1700 x g for 3 minutes. Each sample, was then added an equal volume as the eluate of the lysation buffer AL and 100% ethanol and vortexed. The mixture was transferred to a spin column and the manufactures protocol followed until the elution step where 75 µl EB was used instead of 200 µl. All handling of samples, from sampling water to extraction of eDNA and handling of extracts, was performed under strict clean conditions at designated clean labs for eDNA work at the Norwegian College of Fishery Science.

For the initial round of amplification, each PCR was conducted in a 25µl reaction volume containing 10ng of template DNA, 0.5µM of each primer and 12.5 units of Q5 Hot Start High-Fidelity 2X Master Mix (New England Biolabs: Ipswich, Massachusetts, USA). The following primers were used in the PCR reactions to amplify the 18S V9 region: forward primer 1391_F (5'-GTACACACCGCCCGTC-3') and reverse primer 1560_R (5'-TGATCCTTCTGCAGGTTCACCTAC-3') with Illumina adapter sequences attached in the PCR cycle to amplify sequences for the MiSeq sequencing run. The following PCR profile was then used to amplify fragments for MiSeq sequencing: one initial cycle of fifteen minutes at 95°C; 35 cycles of 45s at 95°C, 45s at 58°C, and 60s at 72°C; and one final cycle of ten minutes at 72°C. The PCR profile of the fragment amplification for Ion Torrent sequencing only differed from this protocol in the duration of the different temperature phases within the 35-cycle loop, which were 30s for each of the three phases. The PCR products were then run on a 2% agarose gel for quality control using SYBR safe (Thermo Fisher Scientific, Waltham, MA, USA) as an in-gel stain at x1 concentration and DNA bands were then visualized for inspection under UV light.

Illumina MiSeq paired-end sequencing of the 18S V9 region was carried out at the Glasgow Polyomics lab (Glasgow, Scotland, UK) using the MiSeq reagent kit (600 cycle) (Illumina, San Diego, CA, USA) and 2 × 300bp sequencing. Ion Torrent sequencing of the amplicons was carried out in the Systems Biology Centre of the University of Plymouth (Plymouth, England, UK) on Ion 318v2 Chips, using the Ion PGM Sequencing 200 Kit v2 (Life Technologies Ltd, Carlsbad, CA, USA) for sequencing of up to 200bp. For both sequencing procedures adapter sequences were trimmed automatically before the sequence data was exported as FASTQ files.

### Sequence processing, OTU clustering and taxonomic assignment

Raw reads from the Illumina MiSeq and the Ion Torrent run were processed in the same manner using identical parameters, apart from some differences in pre-processing of the paired-end reads of the Illumina run. All raw reads, single for the Ion Torrent and both paired ends for the MiSeq run, were trimmed with Sickle version 1.33 (Joshi and Fass, 2011) using a quality threshold of 20 and a minimum length of 100 base pairs. For the paired ends, a file with singletons was created at this step, which was excluded from further analysis. The reads were further trimmed using FASTX-Toolkit version 0.0.14 (Hannon Lab) to a maximum length of 200 base pairs. All sequences were then aligned to a reference consisting of representative 18S sequences of all target organisms or a closely related species (GenBank accession numbers: AF208263.1 (*Lepeophtheirus salmonis*); KT989881.1 (*Neoparamoeba perurans*); KJ756812.1 (*Prymnesium parvum*); JF308618.1 *(Pseudo-nitzschia seriata*); EU478793.1 (*Pseudo-nitzschia delicatissima*) in an attempt to filter out non-18S reads. The alignment was carried out in bowtie2 version 2.2.6 ^12^ using the low stringency local alignment option.

The matched reads then underwent further quality checking using PRINSEQ-lite version 0.20.4 (PRINSEQ, 2013) to identify any formatting errors and remove read headers as well as convert the sequences into FASTA format to facilitate downstream processing. The paired-end MiSeq reads were further processed by merging the mate pairs into one sequence using Velvet version 1.2.09 ^13^ after reversing and complementing reads two. Merged MiSeq and single Ion Torrent reads were further processed using USEARCH version 8.1.1861 ^14^: reads were scanned for unique sequences, sorted and finally clustered into operational taxonomic units (OTUs) using the UPASRSE algorithm and a 97% identity threshold.

A table listing the OTUs and read frequency for every sample was constructed for each of the two the two sequencing methods. Taxonomic identity of the OTUs was assigned in Qiime version 1.9.1 ^15^ using the closed reference approach with the SILVA database release 128 ^16^ as a reference, which is the most comprehensive database for eukaryotic 18S sequences, and the BLAST algorithm for assignment. Multiple identical assignments for different OTUs were pooled together using Primer6 version 6.1.4 ^17^.

Sequence reads generated in this study have been submitted to the NCBI short-reads archive (SRA), accession number XXXX.

### Sequencing efficiency of MiSeq versus Ion Torrent

Sequencing success of MiSeq sequencing compared to Ion Torrent sequencing was quantified by the percentage of raw reads retained after quality filtering as well as by total numbers of unique OTUs identified. Overall sequencing output and sample composition for both sequencing methods were further explored by determining absolute and relative read abundances using the phyloseq package version 1.18.0 ^18^ in the statistical analysis software R version 3.3.0 ^19^. The effect of treatment, sequencing method and medium type on the number of OTUs and evenness index J was tested using General Linear Models of the form:

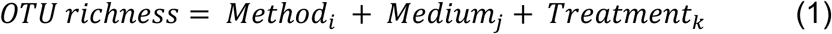

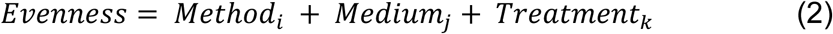

Where: method is a categorical variable with two levels (*i* = Illumina MiSeq, Ion Torrent), Medium is a categorical variable with two levels (j = filtered, unfiltered), Treatment is a categorical variable with 12 levels (*k* = LeSa _2,6,18_, PaPe _3, 6, 18_, AlgMix_3,6,18_, Mix_3,6,18_).

### Sensitivity of MiSeq versus Ion Torrent to reflect sample composition

Potential differences between the two sequencing methods in reflecting OTU composition in the samples was explored using multivariate statistics. For this analysis the OTU data were first to account for inter-sample variation in sequencing depth. Data normalization was conducted in the R package Deseq2 version 1.14.0 ^20^ with which instead of rarefying, a variance stabilizing transformation was applied to the read numbers to convert the counts so that they are of homoscedastic distribution, after the recommendations of ^21^. The normalized read samples were then analyzed within each method for pairwise similarity using the Bray-Curtis similarity index. The pairwise similarities between all samples were then visualized using Multidimensional Scaling Ordination (MDS).

### Sensitivity of MiSeq versus Ion Torrent to reflect the abundance of target species

The two sequencing methods of MiSeq and Ion Torrent were compared in their efficiency in reflecting the true abundances of species within each medium type i.e. filtered and unfiltered medium using a General Linear model of the form:

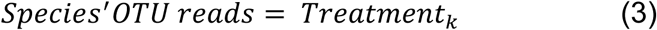

Where k = 2, 6 and 18 abundance levels. The first order interaction between treatment and method and treatment and medium was also tested in order to check whether the effect of treatment depends on which method or which medium is being used. The relative (normalized) reads were used for this analysis and for the unfiltered samples we subtracted background concentrations of our target species that were found in the blank from the abundances found in the treatments.

## Results

### Sequencing efficiency and taxonomic assignment of MiSeq versus Ion Torrent

The Illumina platform successfully sequenced 68 out of the 84 samples whilst Ion Torrent returned a slightly lower number (62 out of the 84 samples). The two sequencing approaches showed differences in their raw read numbers and overall sequence read quality. The Illumina MiSeq data consisted of fewer raw reads (6,115,810 paired reads) across all samples compared with the Ion Torrent output (9,350,400 single reads). After quality filtering of the raw reads, 93.7% of the MiSeq sequence pairs were kept compared to just 68.3% of the Ion Torrent raw reads. As a result the total number of reads that passed QC were similar for both sequencing methods. Samples from the *L. salmonis* incubation in unfiltered samples failed sequence on both platforms, presumably due to problems with the first round PCR.

The OTUs of our algal species of interest were the most abundant reads the filtered samples containing the algal mix, and their identity was validated by a nucleotide BLAST search against the Genbank database on the NCBI website. Both sequencing methods assigned taxonomic identities to our target algae species that were different to their known identity. The OTU that was assigned to *Prymnesium* was first mis-assigned as *P. nemathicium* using SILVA but later correctly identified as *Prymnesium parvum* by the top NCBI BLAST hit. As such this OTU was identified as *P. parvum* for downstream analysis. The validation of the *Pseudo-nitzschia* spp. identities was more ambiguous since our target species, *Pseudo-nitzschia seriata* and *P. delicatissima,* were assigned as *Pseudo-nitzschia australis* and *cuspidata* using SLVA and these taxonomic identities were confirmed by the top BLAST hits for these OTUs. *L. salmonis* and *P. perurans* were correctly assigned to species level with reference to both SILVA and NCBI BLAST.

Once the OTU tables from each method were collapsed to avoid multiple identical taxonomic assignments, the Ion Torrent data appeared to capture slightly more diversity across all samples (2463 OTUs) compared to MiSeq (2277 OTUs). This was also observed on a per treatment basis (Figure 2A, B) as the effect of method was found significant (GLM, F-ratio= 13.5, p<0.001) having accounted for medium type (pre-filtered vs unfiltered seawater). As expected the number of OTUs was much larger in the unfiltered medium type than the pre-filtered samples (GLM, F-ratio= 1329.8, p<0.001). The effect of method depended on the medium type with the Illumina MiSeq performing significantly better than Ion Torrent in detecting more OTUs in the unfiltered sea water medium compared to the filtered one (GLM, F-ratio= 16.5, p<0.001). Sample evenness (i.e. how evenly reads are distributed across OTUs) was significantly lower across treatments (Figure 2C,D) for the Ion Torrent method (GLM, F-ratio = 17.2, p-value<0.001) and this was consistent both in the filtered and the unfiltered medium (GLM, F-ratio = 1.3, p-value=0.249).

**Figure 2:**
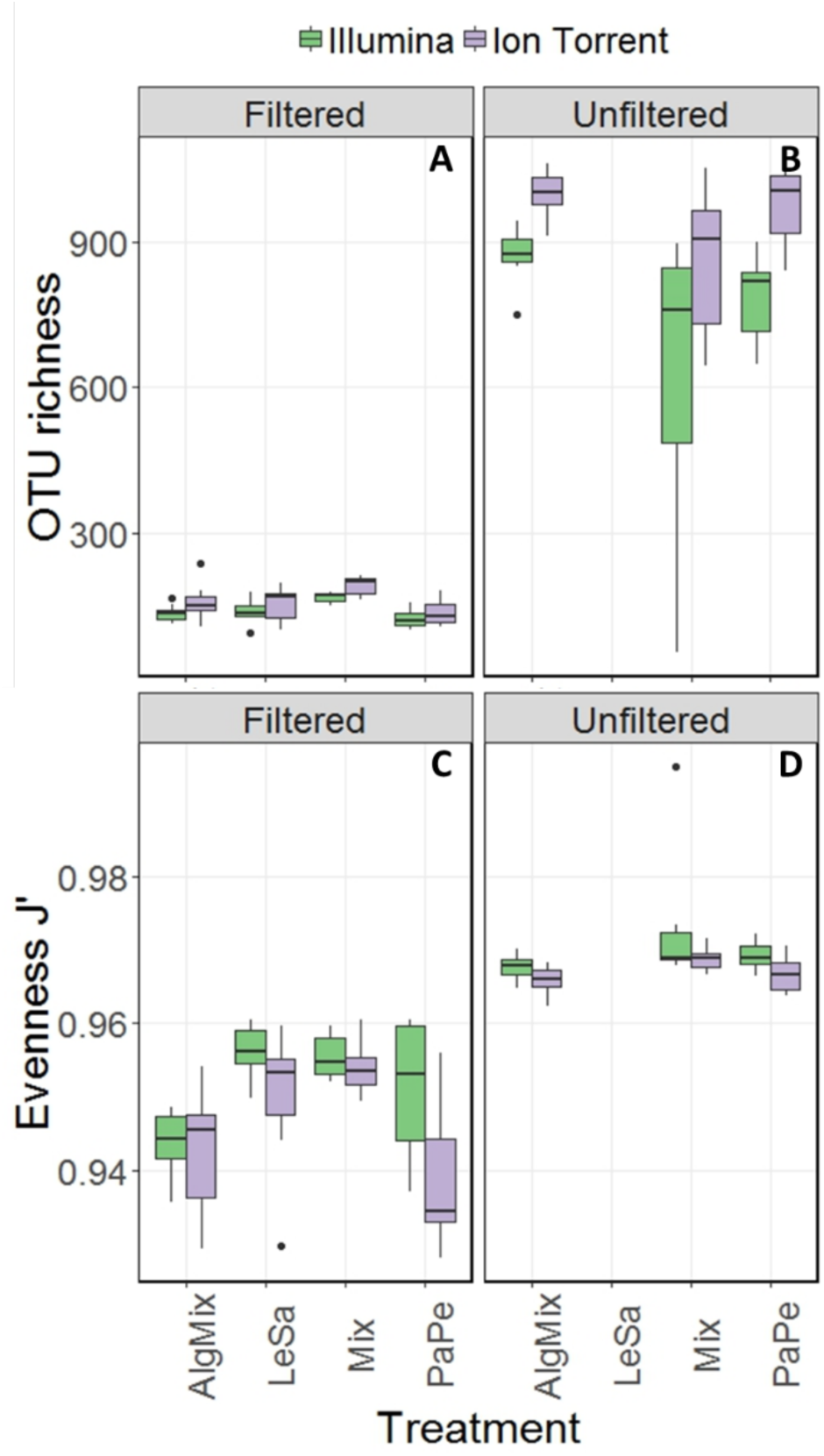
Comparison between Illumina MiSeq (green bars) and Ion Torrent sequencing (grey bars) in reflecting OTU richness (panels A,B) and evenness (panels C,D) across the four treatment levels: *Lepeophtheirus salmonis* (LeSa), *Paramoeba perurans* (PaPe), algal mix (AlgMix) containing the three algal species, and Mix containing all the five species together. Within each panel the two different methods are compared.

### Sensitivity of Illumina MiSeq versus Ion Torrent to reflect sample composition

Both methods detected similar proportions of taxonomic groups considering the absolute read numbers across incubation treatments (Figure 3). Treatments containing unfiltered medium were more diverse in other taxonomic groups including bacteria, archaea and eukaryotes other than our target species. By comparison, in the filtered samples only bacteria were present apart from our target species. The Illumina Miseq and Ion Torrent methods were identical in depicting sample OTU composition as seen both by the relative contribution of taxa they detect (Figure 3) as well as the pairwise similarity between treatments (Figure S1). Specifically, community composition in the unfiltered samples was clearly separated from the filtered samples using data from either sequencing method (Figure S1). Also, samples from different treatments were grouped together at equal similarity within each method, showing more pronounced separation of treatments in the pre-filtered medium that the unfiltered one. The greater similarity observed in the unfiltered treatments was because of the interference of the non-target organisms in the seawater sample that were common to all unfiltered samples.

**Figure 3.**
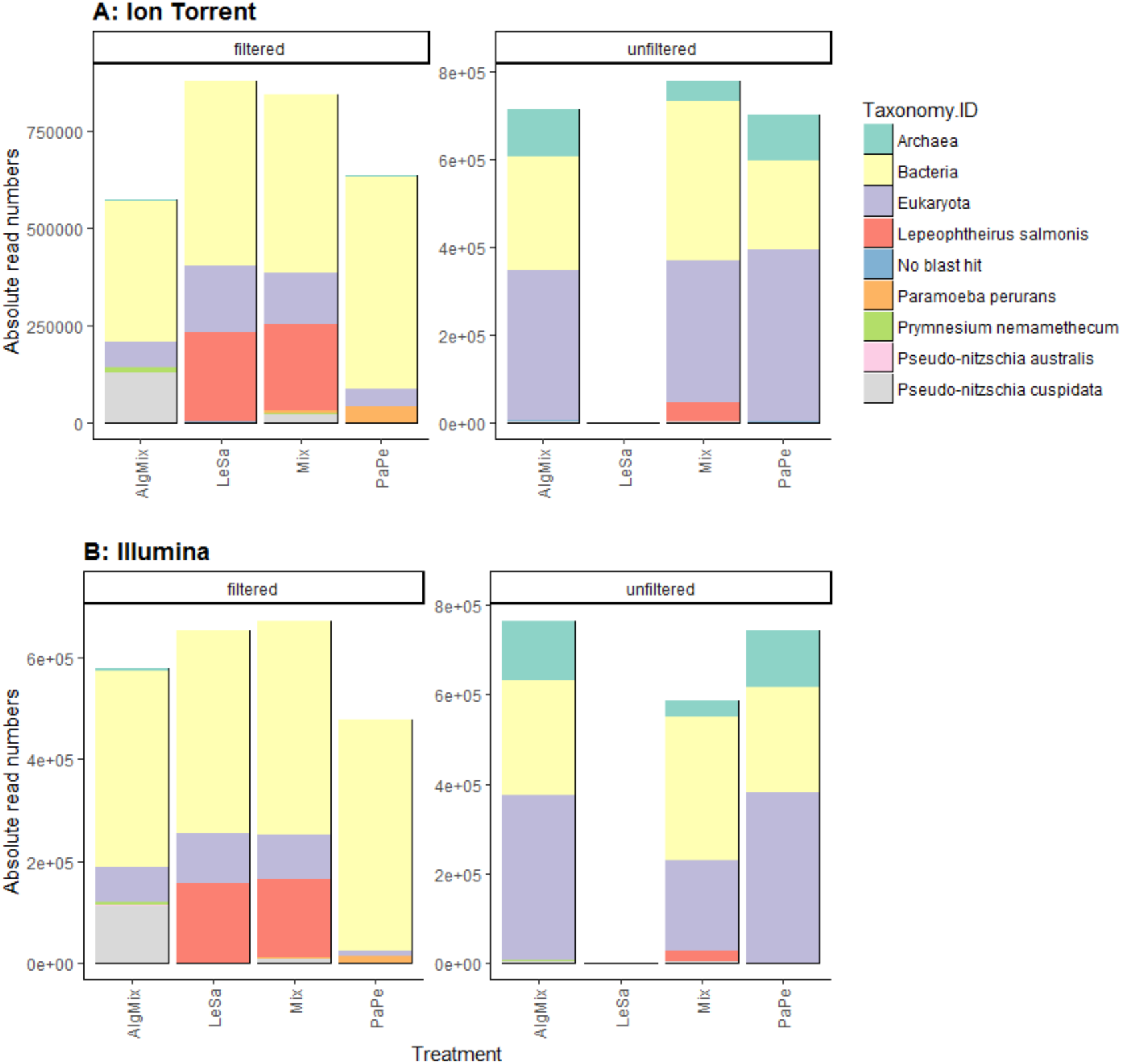
Absolute read numbers for samples grouped according to treatment and split for medium type. Color code identifies different taxonomic groups. Panel A represents the MiSeq data, panel B the data from the Ion Torrent run.

**Figure S1.**
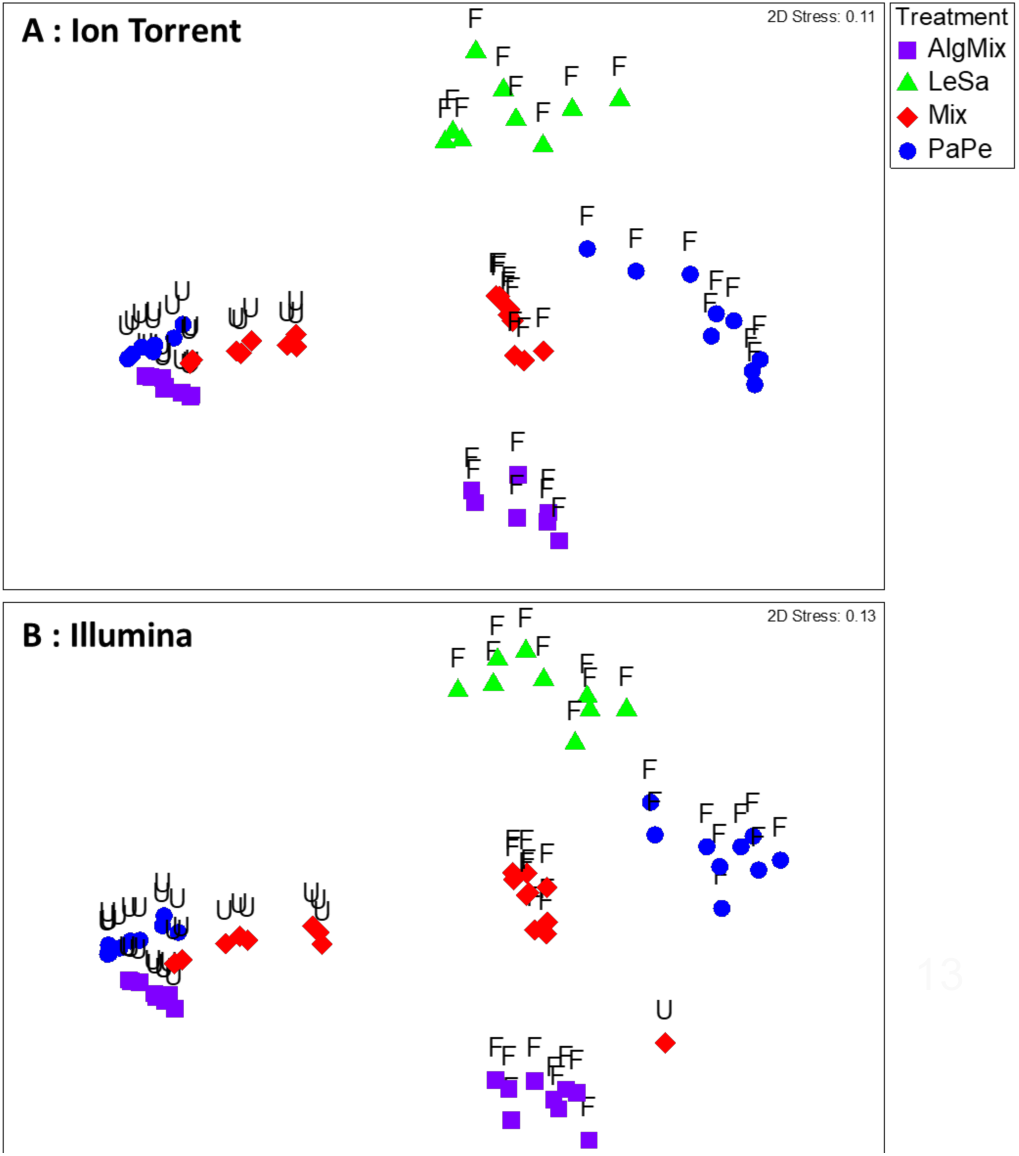
Multidimensional scaling ordination plots showing the similarity between samples based on their OTU-read composition. Samples are color coded for the four treatment levels: *Lepeophtheirus salmoni*s (LeSa), *Paramoeba perurans* (PaPe), algal mix (AlgMix) containing the 3 algal species, and Mix containing all the five species together. Panel A shows the pairwise sample similarity based on data derived from Ion Torrent sequencing, and panel B from Illumina MiSeq sequencing.

### Sensitivity of Illumina MiSeq versus Ion Torrent to reflect the abundance of target species

In the filtered samples, the two sequencing methods were not able to reflect the different abundance levels of the two parasite species *L. salmonis* and *P. perurans* (Figure 4A,C and Table 1) and the same was true for the harmful microalgae species (Figure 5A,C,D and Table 1). In the filtered samples however, the two methods were able to detect the tripling in abundance between the three abundance treatments (2, 6 and 18) for all target species apart from *P. australis* (Figure 4B,D and Figure 5B,D,F and Table 1). Specifically, Ion Torrent was successful in 6 out of the seven sequenced treatments whereas Illumina MiSeq for 5 out of seven treatments (see Table 1).

**Figure 4.**
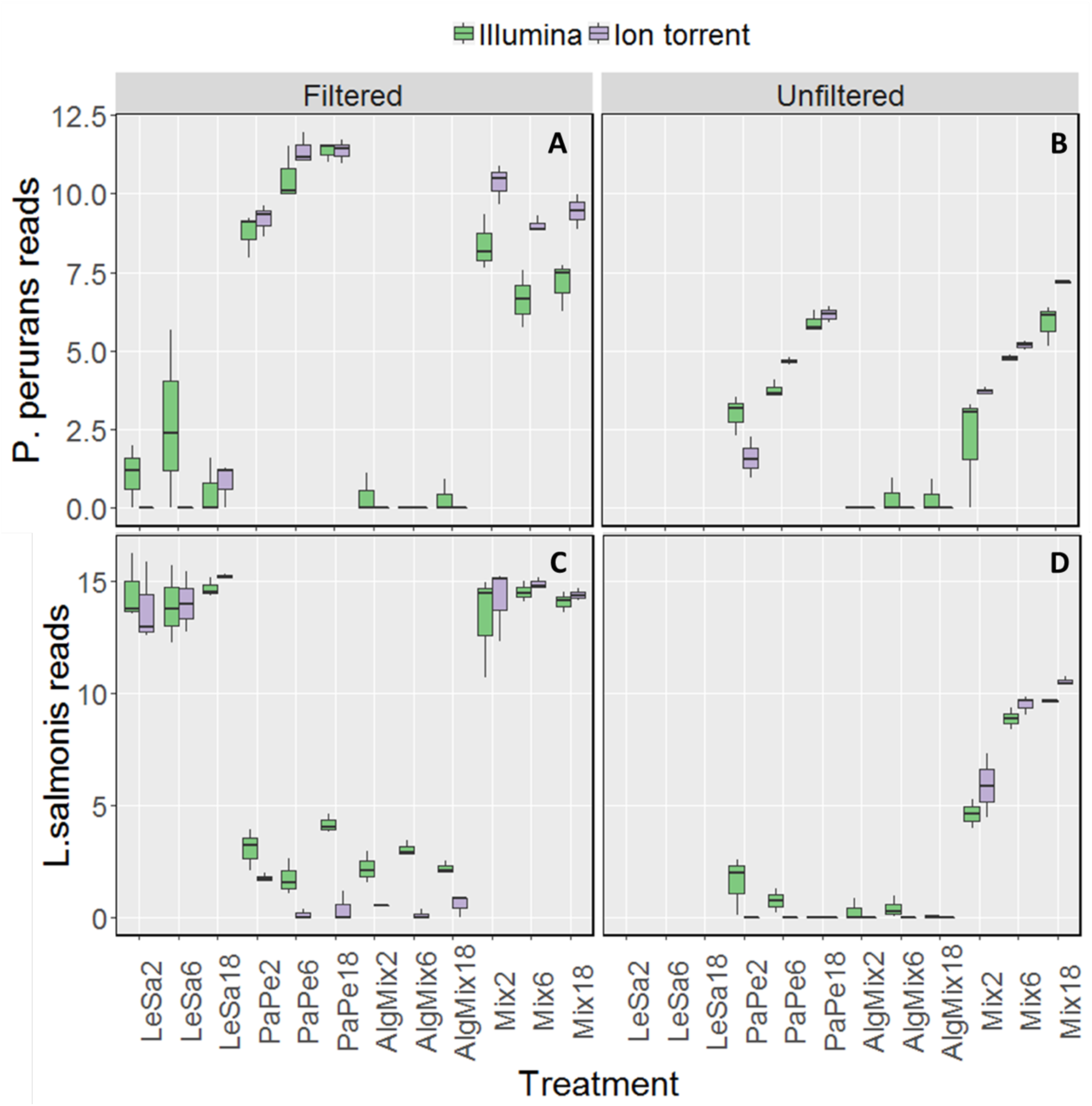
Variability in the read numbers of the two target parasite species namely *Paramoeba perurans* (panels A, B) and *Lepeophtheirus salmonis* (panels C, D) across the different treatments (see figure 1 for treatment abbreviations). Within each treatment, the reads of each species were compared between the two sequencing methods Illumina (green bars) and Ion Torrent (grey bars) and between filtered and unfiltered marine plankton samples.

**Table 1.**
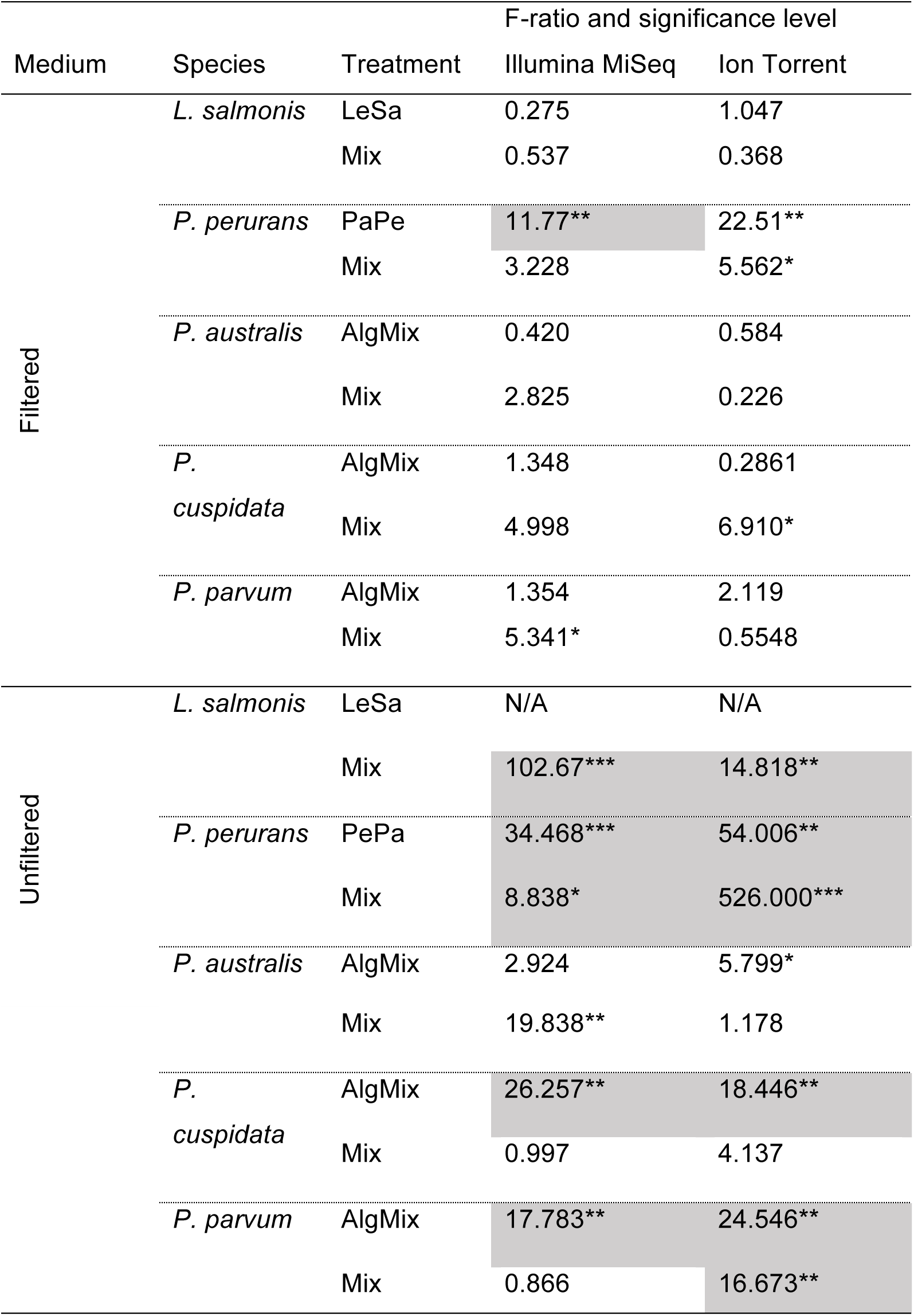
The F-ratio and corresponding confidence level (* for 95%, ** for 99% and *** for 99.9% level) shows whether there is a significant difference between the 3 abundance levels for a given species within the treatment it was present. Dark boxes indicate significant differences and also correct directionality of the effect meaning that an increase in the treatment abundance level corresponded to an increase in the reads that were sequenced.

| Medium | Species | Treatment | F-ratio and significance level |  |
| --- | --- | --- | --- | --- |
|  |  |  | Illumina MiSeq | Ion Torrent |
| Filtered | *L. salmonis* | LeSa | 0.275 | 1.047 |
|  |  | Mix | 0.537 | 0.368 |
|  | *P. perurans* | PaPe | 11.77** | 22.51** |
|  |  | Mix | 3.228 | 5.562* |
|  | *P. australis* | AlgMix | 0.420 | 0.584 |
|  |  | Mix | 2.825 | 0.226 |
|  | *P. cuspidata* | AlgMix | 1.348 | 0.2861 |
|  |  | Mix | 4.998 | 6.910* |
|  | *P. parvum* | AlgMix | 1.354 | 2.119 |
|  |  | Mix | 5.341* | 0.5548 |
| Unfiltered | *L. salmonis* | LeSa | N/A | N/A |
|  |  | Mix | 102.67*** | 14.818** |
|  | *P. perurans* | PePa | 34.468*** | 54.006** |
|  |  | Mix | 8.838* | 526.000*** |
|  | *P. australis* | AlgMix | 2.924 | 5.799* |

|  |  |  |  |
| --- | --- | --- | --- |
|  | Mix | 19.838** | 1.178 |
| *P. cuspidata* | AlgMix | 26.257** | 18.446** |
|  | Mix | 0.997 | 4.137 |
| *P. parvum* | AlgMix | 17.783** | 24.546** |
|  | Mix | 0.866 | 16.673** |

**Figure 5.**
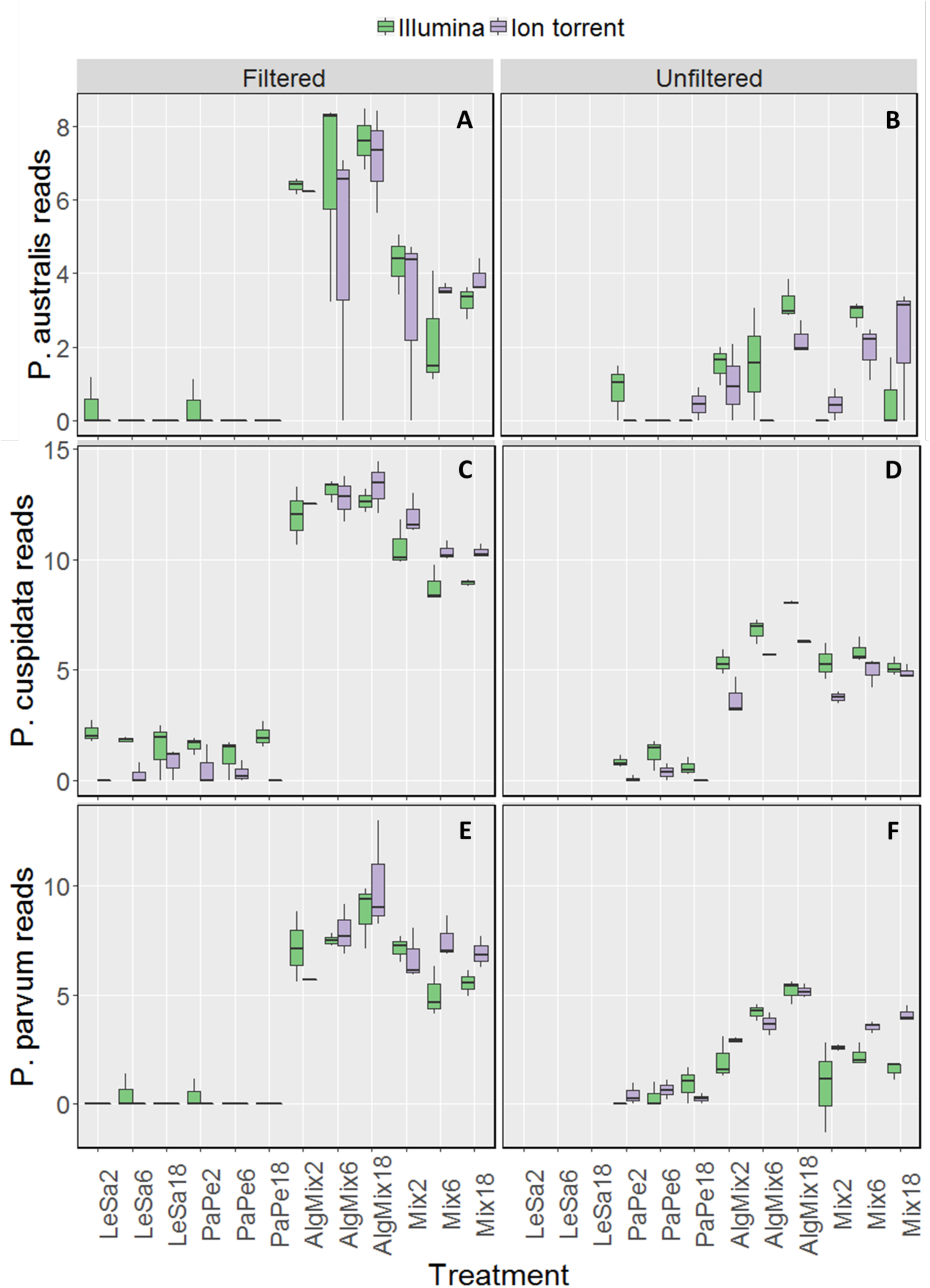
Variability in the read numbers of the three target microalgal species namely the two *Pseudo-nitzschia* species that were assigned as *P. cuspidata* and *P. australis*, but which were in fact *P. delicatissima* and *P. seriata* (panels A, B, C, D) and *Prymnesium parvum* (panels E, F) across the different treatments (see figure 1 for treatment abbreviations). Within each treatment, the reads of each species were compared between the two sequencing methods Illumina MiSeq and Ion Torrent and between filtered and unfiltered marine plankton samples.

## Discussion

The data we present suggest that eDNAseq of the 18S SSU rDNA v9 region can sensitively and specifically identify multiple aquaculture pathogens from the complex mixture of organisms present in seawater. Furthermore, our benchmarking indicates that such detection can be achieved using a low-cost, scalable Ion Torrent sequencing platform – in some cases with better results than the Illumina gold standard. Our findings suggest that eDNAseq may represent a valuable tool in the hands of producers and regulators alike.

As our data suggest, however, several challenges remain. Taxonomic ambiguity around the assignment of the target HAB species (*P. parvum*, *Pseudo-nitchzia* sp.) highlights the difficulty of using short amplicons to assign taxa. Although the 18S V9 SSU region is thought to be useful for detecting global protist diversity ^22^, one size rarely fits all when choosing metabarcoding markers. The 18S SSU rDNA V4 region has also been shown to have high global eukaryotic discriminatory power ^23^. However, to an extent markers must be targeted at particular groups, and the 23S rDNA locus is thought to provide better discrimination for algae in particular ^24^. Nonetheless the 18S v9 region we tested did provide excellent discrimination for *L. salmonis* and *P. perurans* which together account for persistent morbidity in salmonid aquaculture ^9-10^. In future work, it may be more appropriate to target specific organisms with tailored molecular probes, or to deploy amplicon-seq using longer read sequencing technologies to improve global species resolution. However, the former precludes the identification of novel pathogenic agents, while the latter remains too expensive to be widely adopted ^25^.

Species quantification as well as identification remains a holy grail for eDNA studies. Several authors report progress towards this goal in aquatic organisms ^6^, with more recent studies incorporating models of DNA shedding and degradation ^26^. Absolute individual-level quantification is complicated in comparisons of metazoans and unicellular species where biomass, instead of count data, is likely to show better correlation with DNA quantity ^26^. Normalization of read data increases their accurateness in representing actual species abundances, as we have done here, so that experimentally induced differences can be more reliably reflected ^27^. Nonetheless, PCR artefacts relating to runaway amplification of relatively abundant target species templates in pre-filtered water may have negatively impacted relative quantification in our study. However, in more biologically realistic samples (normal, unfiltered seawater) we were able to detect the expected relative abundances according to our dilution series. Our approach can in theory be calibrated to detect absolute abundances – however, PCR always carries an intrinsic risk (e.g. via differential presence of PCR inhibitors in different samples) in quantification from real-world samples ^28^. PCR-free approaches, probe-based, for example, may be more appropriate for the detection of absolute biomass abundance.

Our benchmarking of Ion Torrent and Illumina platforms are in line with the reports of others who have done so for amplicon-seq datasets, especially in relation to higher error rates in the former ^29^. However, both platforms were able to recover community compositions in our data with similar accuracy, as well as recover relative quantities of target species ^29a^. The ability to run a single low output Ion-Torrent chip at a fraction of the price of the Illumina MiSeq to rapidly process only 20-40 samples, means that this technology may be more easy to adopt in a diagnostic context.

Our data, and that of others, does suggest that eDNAseq, may soon become a useful tool for monitoring biological threats to aquaculture. Next steps could include field trials of such methodologies and corroboration with direct count data to fully validate the approach for use in the industry.

## Acknowledgements

This work was supported by the Wellcome Trust [105614/Z/14/Z], a strategic grant from UiT The Arctic University of Norway and BBSRC project BB/N024028/1. Thanks to Catherine Collins of Marine Scotland Science for providing the *P. perurans* and to Nina Lundholm of Natural History Museum of Denmark for providing *P. seriata* used in this study.

